# Generating kinetic environments to study dynamic cellular processes in single cells

**DOI:** 10.1101/632687

**Authors:** Alexander Thiemicke, Hossein Jashnsaz, Guoliang Li, Gregor Neuert

**Author notes:** equally contributing (Alexander Thiemicke, Hossein Jashnsaz). corresponding author: Gregor Neuert.

## Abstract

Cells of any organism are consistently exposed to changes over time in their environment. The kinetics by which these changes occur are critical for the cellular response and fate decision. It is therefore important to control the temporal changes of extracellular stimuli precisely to understand biological mechanisms in a quantitative manner. Most current cell culture and biochemical studies focus on instant changes in the environment and therefore neglect the importance of kinetic environments. To address these shortcomings, we developed two experimental methodologies to precisely control the environment of single cells. These methodologies are compatible with standard biochemistry, molecular, cell and quantitative biology assays. We demonstrate applicability by obtaining time series and time point measurements in both live and fixed cells. We demonstrate the feasibility of the methodology in yeast and mammalian cell culture in combination with widely used assays such as flow cytometry, time-lapse microscopy and single-molecule RNA Fluorescent *in-situ* Hybridization. Our experimental methodologies are easy to implement in most laboratory settings and allows the study of kinetic environments in a wide range of assays and different cell culture conditions.

## Introduction

In a human body, cells are constantly exposed to diverse physiological environments that change over time and space. For example, it has long been known that external or internal stressors^1–4^, morphogen concentrations^5,6^, drugs (pharmacokinetics) or hormone concentrations^7–9^ change over time (Figure 1a, b). Therefore, cells must have developed mechanisms to integrate kinetic changes as well as spatial gradients in the environment and respond to these in a manner benefitting the organism.

**Figure 1:**
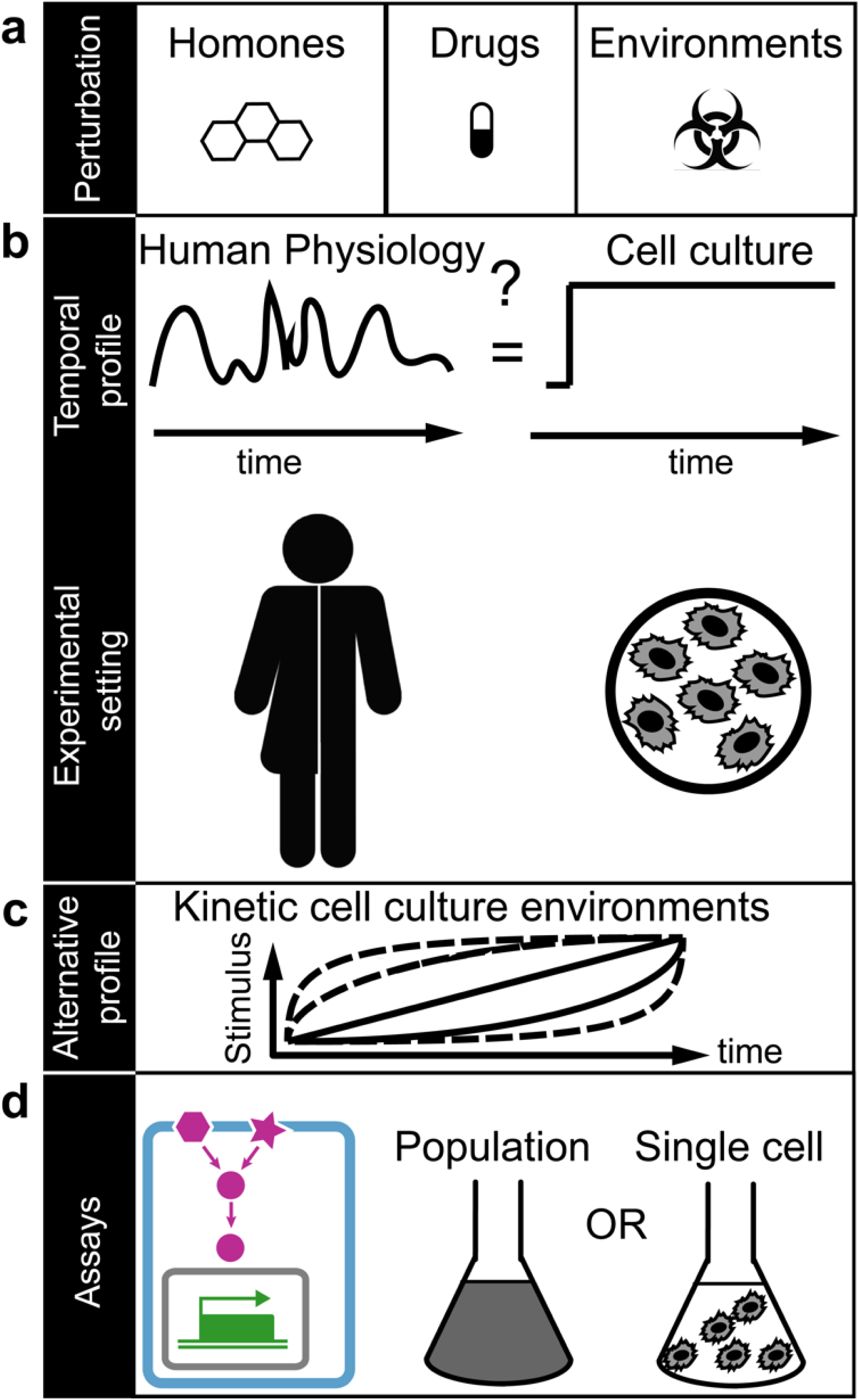
Kinetic cell culture environments mimic physiologically relevant cell environments. (**a**) Examples of different types of cellular environmental perturbations. (**b**) Temporal profile of physiological relevant environment that may fluctuate over time as experienced by cells in humans (left). In contrast, the majority of cell culture-based experiments are performed in constant environments over time and may neglect physiologically relevant conditions (right). (**c**) We propose alternative profiles to study cells in precisely controlled kinetic cell culture environments. (**d**) The power of this approach is demonstrated on measurements of single cell volume changes (cyan), changes in signal transduction (magenta) and gene regulation (green) in cell population (middle) and single cell experiments (right) in yeast and human cells.

Precisely how a single cell integrates kinetic changes in the environment is often not understood. Many previous and current biomedical studies have focused on how cells are affected by sudden or instant kinetic changes of the environment (Figure 1b). The underlying assumption made in these types of experiments is that rapid changes in the environment serve as an adequate representation of a given physiologic or pathophysiologic cellular environment (Figure 1b). For example, physiologic changes in concentrations over time of stresses^1–4^, drugs or hormone levels^7–9^ may be drastically different from instant changes and may result in a different cellular response (Figure 1b).

Recent pioneering studies have provided insights into how molecular processes differ in kinetic environments in comparison to instant changes in the environment^2–5,10–15^. These studies have often relied on two main approaches to deliver the desired kinetic perturbations on the cells. One is the use of specialized, small-scale and custom designed microfluidic setups ^16–20^. But fabricating these devices is often complex, may require specialized equipment to produce, may take significant time to set up and are limited in the compatibility with and application of many common biological assays^16–19^. On the other hand, simpler approaches implemented the use of syringe pumps and flow chambers that avoid the complexity of microfluidics^21^. Although these are state of the art methods to observe single cells in small volumes under the microscope, these methods are not compatible with standard molecular and cell biology assays. Furthermore, currently published methods generate only an approximate kinetic perturbation while they lack in validation and detailed description in how to generate kinetic environments when concentration and volumes changes over time. Therefore, a simple and precise methodology of generating a variety of kinetic environments in regular and microfluidic cell culture do not exit.

Here we provide a simple, precise and versatile methodology to generate a variety of kinetic perturbations for many common bulk and single cell assays by avoiding the complexity of common microfluidic devices. To the best of our knowledge there is no other method yet that enables the study of cells with many biological assays after exposure to a kinetic environment. We describe two methodologies to generate gradually changing environments that are compatible with standard molecular biology and cell culture conditions (Figures 1–3). We demonstrate our methodologies in yeast and a human cell line and demonstrate its applicability in combination with standard laboratory assays of signal transduction and gene regulation in population or single cell experiments (Figures 1d, 4). We believe our simple yet general methodology to precisely control the environment will enable new biological insights in how single cells respond to kinetic changes in the environment.

**Figure 2:**
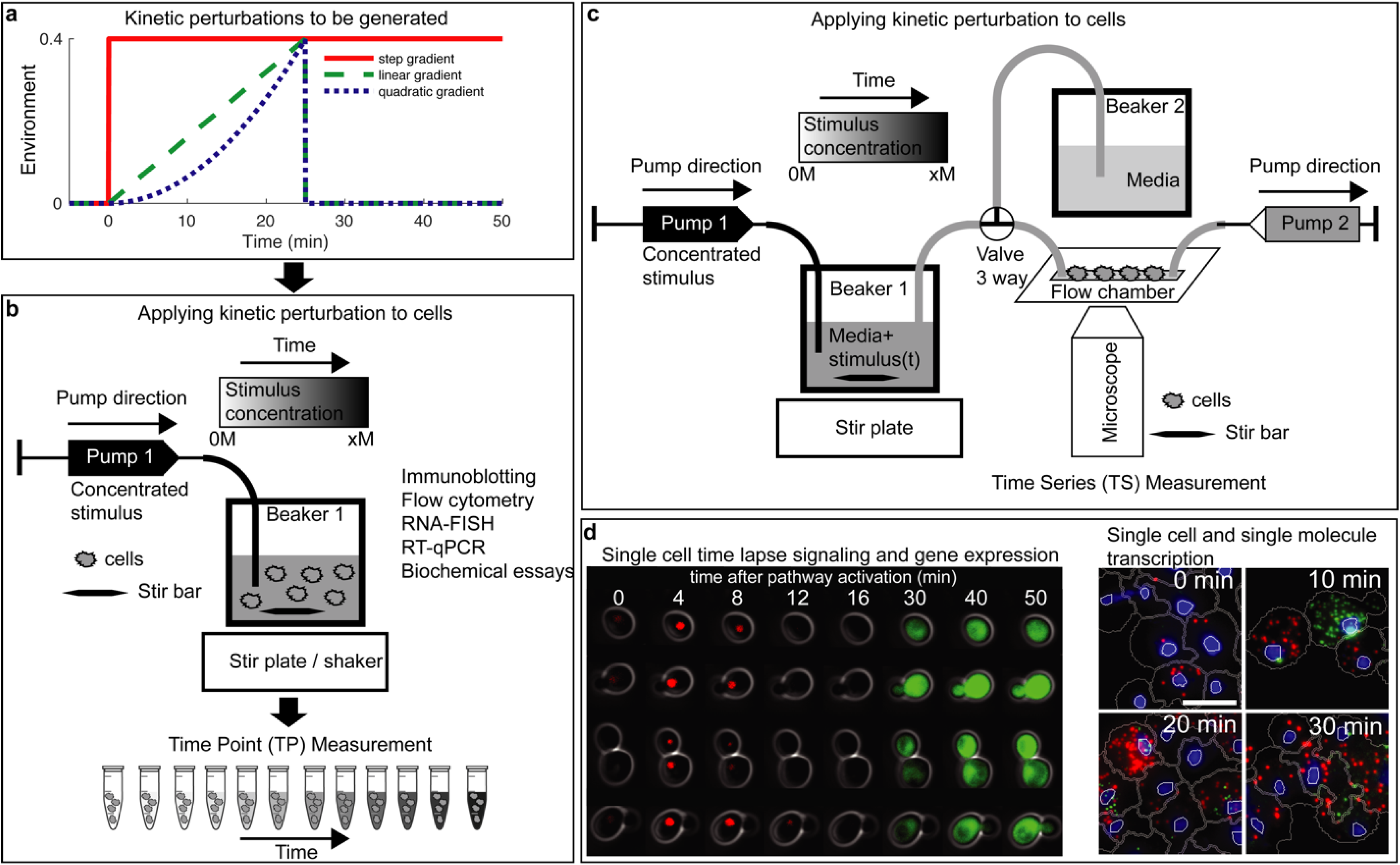
Experimental setup to generate kinetic environmental profiles in the laboratory. (**a**) Three possible kinetic environmental profiles: Step (red), linear gradient (green) and quadratic gradient (blue). (**b**) Experimental setup to generate temporal gradients and measure cells at specific time points. A high concentration of stimulus is added through Pump 1 at constant stirring condition and samples are collected at predefined time points and sample volumes (Time Point (TP) data collection). (**c**) Experimental setup to follow single cells over time in a microfluidic flow chamber on a microscope. Cells are exposed to normal media (Beaker 2) or to a temporal gradient generated in Beaker 1. Pump 1 adds a concentrated substance to Beaker 1 at constant stirring. Pump 2 allows to adjust the flow rate. Time series (TS) data collection is done by tracking cells in the flow chamber using bright field and fluorescent microscopy. (**d**) Application images of single cell signal transduction and gene regulation measured over time using the Time Series (TS) data collection protocol (left) and of single cell transcription snapshots measured using the Time Point (TP) data collection protocol (right).

**Figure 3:**
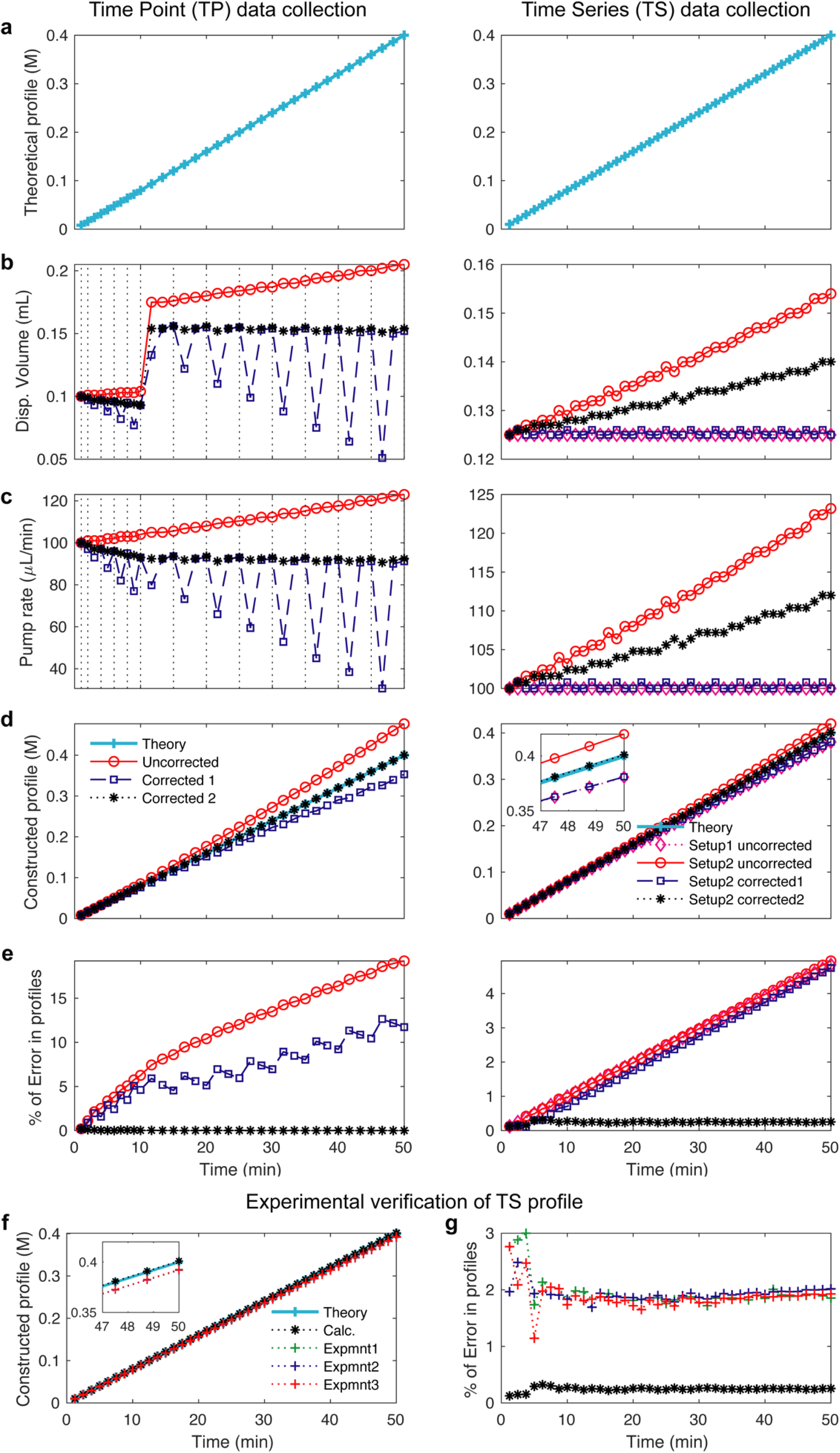
Calculated and experimentally verified pump profiles. (**a-e**) Calculation of pump profile generation for Time Point (TP, left) and Time series (TS, right) data collection. In the TP experiment (left), pump profiles are uncorrected (red), corrected for volume removal during sampling (blue), corrected for volume removal and therefore change in stimulus concentration (black). In the TS experiment (right), two pumps are used. In Setup 1 mixing flask volume (V0) is kept constant by setting Pump 2 rate equal to that of Pump 1 (magenta). In Setup 2, the pump rate for Pump 1 is uncorrected and the pump rate for Pump 2 is kept constant (red). In Setup 2 - correction 1, the pump profile is corrected for volume taken out by Pump 2 (blue). In Setup2 – correction 2, the pump profile is corrected for volume removal and therefore change in stimulus concentration (black). (**a**) Proposed linear pump concentration profile of 0.4 M NaCl. **(b)** Computed and instrument adapted syringe dispense volume. (**c**) Computed pump rate profile over time. (**d**) Computed concentration profiles over time. (**e**) Error comparisons between pump profiles. (**f-g**) Experimentally verified pump profiles. (**f**) The delivered molarities constructed from cumulative dispensed volume measurements via Pump 1 using the setup in Figure 1c in three experiments (red, green, blue) compared to the calculations (black) and theory (cyan). Insert comparing different profiles. (**g**) The corresponding errors from experiments and the calculations compared to the theory.

**Figure 4:**
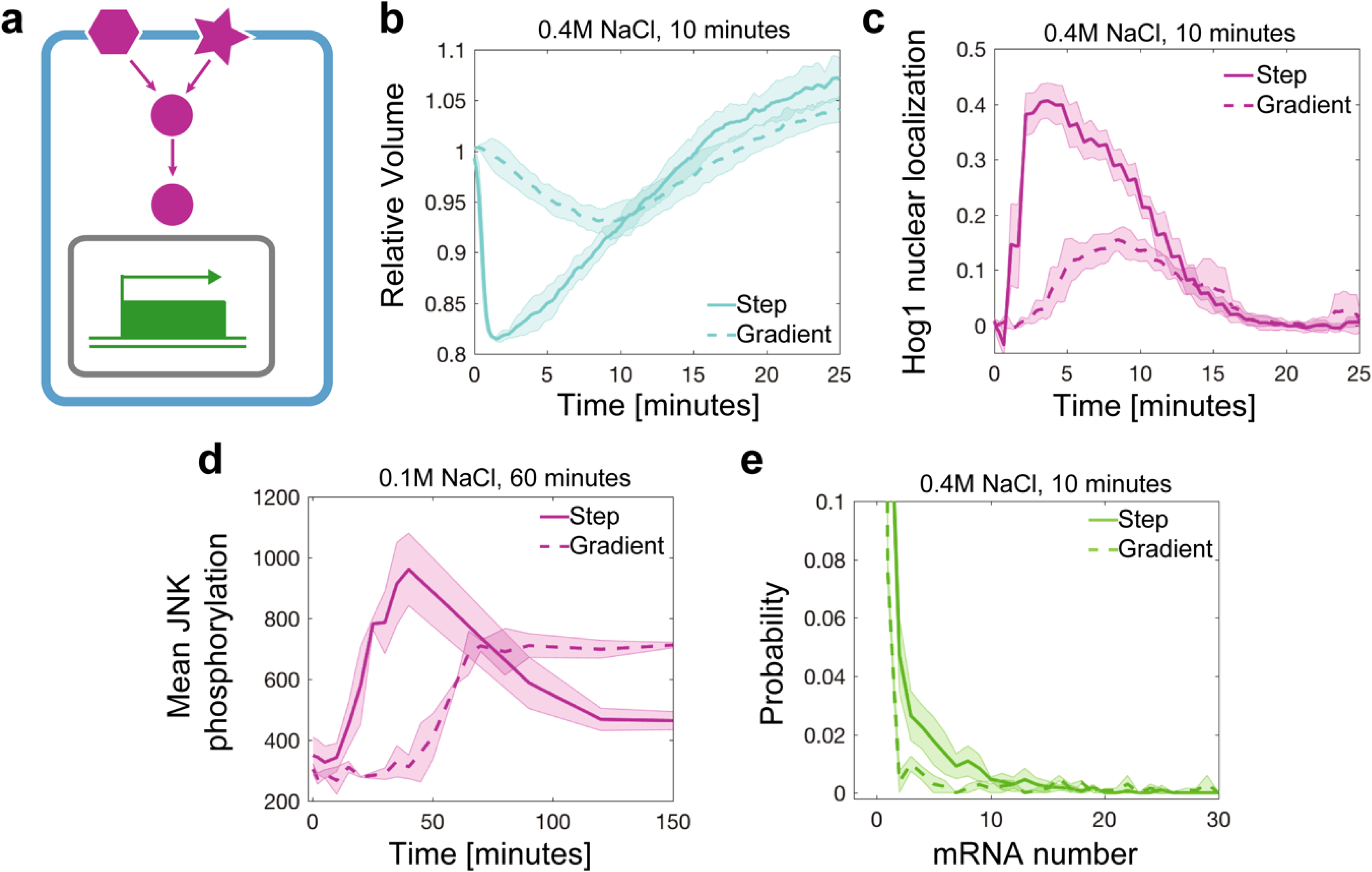
Application of different pump profiles and quantitative assays. (**a**) Overview of cell processes that have been investigated in kinetic environments. (**b**) Quantifying volume and (**c**) signal transduction of Hog1 nuclear localization over time in live single *S. cerevisiae* yeast cells exposed to instant increase to 0.4 M NaCl (dashed line, 79 cells) or to linear gradient of 0.4 M NaCl in 10 minutes (solid line, 90 cells). (**d**) JNK phosphorylation over time measured with flow cytometry in human THP1 cells after exposure to instant increase to 0.1 M NaCl (solid line, 636,628 cells) or to linear gradient of 0.1 M in 60 minutes (dashed line, 1,599,923 cells). (**e**) Single cell distributions of single-molecule RNA FISH measurements of *STL1* mRNA in *S. cerevisiae* yeast cells exposed to instant increase to 0.4 M NaCl (solid line, 3269 cells) or a linear gradient of 0.4 M in 10 minutes (dashed line, 2164 cells). Thick lines are the mean and shaded area are the standard deviation from two or three biological replica experiments single cells.

## Results

### Proof of feasibility using cell lines

To demonstrate our approach, we compare rapidly changing environments to gradually changing environments of increasing concentrations of a stimulus (Figures 1c, 2a). We use osmotic stress to activate the high osmolarity glycerol (HOG) Mitogen Activated Protein Kinase (MAPK) pathway in *S. cerevisiae* yeast cells and the c-Jun N-terminal kinase (JNK) MAPK pathway in a human monocytic cell line. We chose different NaCl concentrations to represent a change in the cellular environment. NaCl is a well-studied stressor in yeast cells and is relevant in human cells in the context of immune cell activation^22–26^ and cell death^13^. We have developed two cell culture methodologies for time point (TP) measurements from continuously growing cells (Figure 2b) or time series (TS) measurements on the same cells over time in a simple microfluidic chamber (Figure 2c). Both methodologies require the use of syringe pumps. We have developed software to accurately compute the pump profiles for a desired experimental design and validated these profiles experimentally (Figure 3). Finally, we demonstrate the feasibility of our methodology on live-cell time-lapse microscopy experiments of cell volume change over time (Figure 4b), dynamic changes in signal transduction in single cells over time (Figures 2d (left), 4c), dynamic changes in protein phosphorylation in human cells using phospho-specific flow cytometry (Figure 4d) and single-molecule RNA fluorescent *in-situ* hybridization quantification of transcription in single cells (Figures 2d (right), 4e).

### Computational pipeline to generate the pump profiles

Concentrated stimulus is added over time to a flask containing media and samples are taken out of the flask for time point (TP) measurements or media is removed in time series (TS) experiments resulting in changes over time of the concentration and volumes in the mixing flask. These changes need to be considered to accurately compute the desired pump profile and failure to do so can result in significant error in the pump profile as plotted in Figure 3. The desired concentration profile consists of a maximum number of discrete time points set by the programmable pump. We construct any arbitrarily concentration profile by combining several short segments with linear concentration profiles. From the beginning of each interval to the end of that interval we increase the concentration linearly with a fixed rate *dr*_*i*_ as shown in Figure S1. However, the rate from each phase to the next could be changed to produce any arbitrary profile over the whole treatment time (*dr*_*i*_=1 *dr*_*i*+1_). During each interval, stimulus over time is delivered continually by adding appropriate amount (*dv*_*i*_) of concentrated stimulus (*C*_*max*_) to the total volume of growth media in a flask. The pump profile is then computed using the following considerations:

1. During each time interval, a defined volume of concentrated stimulus is being added to the total flask volume using Pump 1.
2. At each time intervals, a fixed volume is taken out of the flask. In the case of TP experiment, we sample a fixed volume of cells (Figure 2b) or in case of TS experiments Pump 2 delivers media with a defined stimulus concentration to cells in a flow chamber (Figure 2c).

Removing volume and adding stimulus to the flask result in a concentration change that needs to be accounted for in the pump profile calculation, which is not done by any other published method. In Figure 3 we demonstrate that not taking these considerations into account result in large errors specifically at high stimulus concentrations.

### Algorithm to compute pump profiles

We calculate the stimulus concentration profile for discrete time points as depicted in Figure S1. First, we calculate the theoretical values of any given stimulus concentration profile, *m*(*t*), at a fixed number of time points, [*t*_1_, *t*_2_, *t*_3_,… *t*_*N*_], with time intervals [*dt*_1_, *dt*_2_, *dt*_3_,… *dt*_*N*_]. The time intervals could be chosen either uniformly or variable. We increase the stimulus concentration *m*_*i*−1_ linearly from the beginning of the *i*^th^ interval at *t*_*i*−1_ to *m*_*i*_ at the end of the *i*^*th*^ interval at *t*_*i*_. Pump 1 adds fixed volume *dv*_*i*_ of concentrated stimulus to the mixing Beaker1 during interval *dt*_*i*_ at a fixed pump rate of *k*_*i*_ = *dv*_*i*_ */dt*_*i*_. The beaker has an initial volume of *V*_0_ and an initial stimulus concentration of *m*_0_ = 0 at *t* = 0. The stimulus concentration profile at any given time point (*m*_*i*_) is then calculated as:

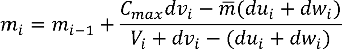

where *C*_*max*_ is the concentrated stimulus (in mM),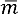 is the average of *m*_*i*_ and *m*_*i*−1_ (in mM), and *dv*_*i*_ (in mL) is the dispensed volume of concentrated stimulus during the time interval *dt*_*i*_. *du*_*i*_ (in mL) is the volume taken out by Pump2 (in TS experiment), and *dw*_*i*_ (in mL) is the volume taken out due to sampling (in TP experiments), both during the interval *dt*_*i*_. Finally, *V*_*i*_ is the total flask volume (in mL) at *t*_*i*_. Once we computed *dv*_*i*_, then we compute the pump rate as *k*_*i*_= 1000\**dv*_*i*_ / *dt*_*i*_ in µL/min. We operate Pump2 at a fixed rate of 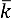, therefore 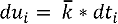, for TS experiments, while we don’t need Pump2 for TP experiment and *du*_*i*_ = 0. We round the calculated values of *dv*_*i*_ in the specified unit to 3 digits after the decimal which is the functional value for the syringe pumps. This calculation is what we refer to setup2 in Figure 3. In setup1, the desired profiles are calculated by setting Pump2 rate equal to that of Pump1 over the treatment duration, which results even in larger errors in the generated profiles. Examples of corrected and uncorrected concentration profiles are shown in Figure 3. Our methodology, once corrected for the volume and concentration changes properly, generates stimulus profiles within 1% error of the theoretical desired profiles (Figure 3).

The profiles are generated under the following conditions:

a. The concentrated stimulus concentration *C*_*max*_ = 4 *M*.
b. The total flask volume *V*_0_ = 50 *mL* at t = 0.
c. Pump2 rate was set to 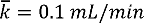 *mL/min* for TS and 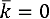 for TP experiment.
d. Samples taken out at the fixed volumes of *dw*_*i*_ = 1 *mL* at the time points [1,2,4,6,8,10,15,20,25,30,35,40,45,50] minutes for TP, while no sampling done for TS.
e. Both TP and TS profiles are generated over 50 minutes. TS in 40 intervals and TP profile in 34 intervals set optimally by the programmable syringe pump.

The calculation results are shown in Tables S1 and S2 for TS and TP profiles.

### Experimental validation of pump profiles

We experimentally verify the specific profile presented in Figure 3 (right, linear gradient of 0.4 M over 50 min for TS experiment). In order to illustrate the high accuracy and precision of the profiles applied to the cells, we experimentally instrument-proof our calculations. We measure the total dispense volume via Pump 1 delivered using the setup in Figure 1c by measuring the weight of the Beaker 1 over the treatment time on a digital balance. As shown in Figure 3f, we calculate the molarity profiles resulting from these measurements and compare them to our calculation and theoretical profiles. In Figure 3g, we show the corresponding errors from experiments and the calculations compared to the theory. These results (errors within 2% of the theoretical values and standard deviation out of 3 experiments is below 1%) show that our setup applies the calculated profiles to the cells with high accuracy and precision. Importantly, these corrections ensure that the pump profiles are accurate over the full-time courses correcting for any nonlinear artefacts as validated experimentally (Figure 3e).

### Time Point (TP) Measurements for cell population and single cell experiments

In the Time Point (TP) Measurement setup, cells are grown in cell culture flasks in a cell culture incubator (Figure 2b). The concentrated stimulus is pumped into the flask through the computer programmable syringe Pump 1 (Figure 2b). In case of the experiments with microbial cultures, cells are mixed with a magnetic stir bar. The mixing speed is optimized to ensure fast mixing of the continuously added concentrated stimulus and at the same time cause little perturbation to the cells. In case of the human cell experiments and some of the yeast experiments, cell culture flasks sit on an orbital shaker that mixes fast enough to ensure rapid mixing but does not interfere with cell growth. At a number of predefined time points, a fixed volume of cells is removed manually with a syringe and collected for further analysis in cell population or single cell assays (Figures 2b, 4d, 4e). For each experiment, the dispensed volume, the number and volume of samples for each time point is defined before the experiment. Based on these experimental parameters, the pump profile is calculated and the pump is programmed. Several considerations are important to ensure correct generation of stimulus profiles (Figure 3): First, the pump program must adjust for the reduction in volume in the flask during sample removal. Second, the pump rate has to decrease over time due to an increase in volume caused by continuous addition of concentrated stimulus. Figure 3a (left) presents an example in which cells are exposed to a linear gradient for 50 minutes to a final concentration of 0.4 M NaCl. When cells are sampled from the cell culture flask for downstream assays (dashed black lines), the cell culture volume and the total amount of NaCl in the culture changes. To better illustrate this point, we computationally compared three possible pump profiles. These are uncorrected pump profiles (red), pump profiles corrected for volume removal during sampling (blue), and the correct pump profiles which are corrected for volume removal and therefore change in total NaCl amount in the flask (black). For each of these cases, we compared the dispensed volume of highly concentrated NaCl solution in ml (Figure 3b), the pump rate in µL per minute (Figure 3c), the computed concentrated profiles (Figure 3d) and the percent of error for each condition (Figure 3e). Initially, the corrections are small, because cells are sampled rapidly within the first 10 minutes. But if the sampling time is not frequent, then corrections have a significant impact on the dispensed volume (Figure 3b) and the pump rate (Figure 3c). The result is that as time progresses, the corrections have a significant nonlinear effect over time on the accuracy of the pump profile (Figure 3d, e).

### Time Series (TS) Measurements in single cell time-lapse microscopy experiments

In the Time Series (TS) Measurements, cells can be grown in a microfluidic flow chamber and simultaneously imaged on an inverted microscope (Figure 2c). This experimental setup consists of syringe Pump 1 that pumps concentrated stimulus into Beaker 1 containing growth media and a magnetic stir bar for rapid mixing (Figure 2c). Connected to Beaker 1 is the microfluidic flowcell and a syringe Pump 2 that pulls liquid from Beaker 1 and over the cells in the flow chamber, generating a temporal gradient of the stimulus. The cells are adhered to the concanavalin A-coated coverslip in the flow chamber, allowing for rapid media exchange over time. To better illustrate how the two pumps work in concert, we simulate the addition of a linear gradient for 50 minutes to a final concentration of 0.4 M NaCl (Figure 3a (right)). Because Pump 2 removes media over time, the volume and concentration changes in Beaker 1 need to be accounted for. Instead of experimentally testing these effects, we first computationally compared four possible pump profiles which are Setup1 – uncorrected in which the mixing flask volume (V0) in Beaker 1 is kept constant by setting pump rate of Pump 2 equal to that of Pump 1 (magenta). In Setup2 – uncorrected, the pump rate for Pump 1 is uncorrected and the pump rate for Pump 2 is kept constant (red). In Setup 2 - correction 1, the pump profile is corrected for volume taken out by Pump 2 (blue). In Setup2 – correction 2, the pump profile is corrected for volume removal and therefore change in stimulus concentration (black). For each setup, we compare the dispensed volume of highly concentrated NaCl solution in ml (Figure 3b), the pump rate in µL per minute (Figure 3c), the computed concentration profiles (Figure 3d) and the percent of error after for each condition (Figure 3e). As time progresses, the corrections become more significant for the dispensed volume and the pump rate for setup 2 uncorrected and setup 2 with correction 2. These differences can be seen in the constructed profiles (Figure 3d). From these computational profiles, it became apparent that the corrections are important as they reduce errors in the pump profiles significantly (Figure 3e).

### Application of kinetic cell perturbation to study dynamic cell shape, cell signaling and gene regulation in single and populations of cells

To demonstrate the power of our approach, which is the ability to precisely control the kinetic environment of a cell, we focused on three levels of cellular response: cell shape, signal transduction and gene regulation (Figures 1, 4a). As physiologically relevant environment, we chose different kinetic gradients of osmotic stress and applied these to *S. cerevisiae* yeast cells and the human monocytic THP1 cell line. In yeast, the High Osmolarity Glycerol (HOG) pathway belongs to the class of Mitogen Activated Protein Kinase (MAPK) that enables cells to respond to changes in external osmolarity^27^. The terminal kinase, Hog1, is evolutionary conserved between yeast and human and is a functional ortholog kinase to JNK in human cells^28^. In humans, JNK is involved in many pathophysiological conditions^29,30^, and JNK can rescue Hog1 function in yeast cells^28^. Upon osmotic stress, yeast cells are osmotically compressed resulting in significant volume decrease as measured in our time series (TS) measurements (Figure 4b). In comparison to instantly changing environments where volume decreases rapidly^2^, slowly changing environments such as a linear increase to 0.4 M NaCl in 10 minutes result in slowly changing volume (Figure 4b). Simultaneously, we measured Hog1 nuclear localization under the same conditions resulting in maximum signaling at 8 minutes after linear osmotic stress in comparison to a maximum in Hog1 nuclear localization after 2 minutes upon an instant change in osmolarity (Figure 4c). Next, we tested our Time Point (TP) measurement protocol on cells that are sampled at different time points after osmotic stress. We exposed human THP1 suspension cells to instant increase of 0.1 M NaCl and a linear gradient increase of 0.1 M NaCl in 60 minutes. We sampled cells between 0 and 150 minutes in intervals of 2, 5, 10, or 30 minutes and fixed cells with formaldehyde. Cells are subsequently permeabilized and then stained with an antibody for phosphorylated form of JNK^31^. From single cell distributions, the average JNK phosphorylation level were computed for three independent biological replica experiments (Figure 4d). Finally, we applied Time Point (TP) measurement protocol to *S. cerevisiae* yeast cells to measure mRNA expression of the osmotic stress response gene *STL1* in single cells. We exposed cells to an instant increase of 0.4 M NaCl and a linear gradient with a final concentration of 0.4 M NaCl in 10 minutes. Figure 4e depicts single cell distributions of single-molecule RNA FISH data 6 minutes after NaCl stress.

## Discussion

We describe methodologies that allows the generation of physiologically relevant environmental changes in cell culture. We demonstrate in Figure 3 that it is critical to consider the specific sampling time, sampling volume, number of samples and volume change caused by the addition of stimulus. Because sampling changes the total number of cells in the flask, the total stimulus and the volume, these conditions need to be reflected in the programed pump profiles. Failure to do so result in wrong pump profiles with non-linear errors over time. We demonstrate in several examples that we generate gradual changes in the environment over time. Our methodology aids in the application of profiles that might more closely represent physiological conditions than rapidly changing environments (Figure 1). We developed two experimental methodologies to measure cells at specific time points, labeled Time Point (TP) and Time series (TS) measurements, that can be combined with standard cell population experiments and single-cell experiments (Figure 2). In contrast to the vast amount of microfluidic approaches that are available, our methodology allows a quick and easy setup that can be established in any biological laboratory setting. Further, because our method is scalable to large volumes of cell culture, it can be combined with microfluidic/live cell microscopy, gene expression analysis, flow cytometry and many other biological and biochemical assays. From these experiments we observe that cell volume and Hog1 signaling changes slowly if the kinetic osmolyte profiles changes rapidly. This is consistent with previous reports on Hog1 signaling during osmotic stress^20^. Mean JNK phosphorylation is delayed and non-adaptive in cells exposed to a gradient in comparison to a step of NaCl. This result indicates that the JNK activation dynamics respond is NaCl threshold dependent. Finally, RNA-FISH data in yeast indicate that *STL1* mRNA expression is delayed if cells are exposed to gradual changing environments.

## Conclusion

We developed algorithms that generate precise kinetic perturbations by accounting and correcting for the changes in cell culture volume and concentration profiles caused by the stimulus application and sample removal. We demonstrate the wide applicability of this approach to study cell response in terms of changes in cell volume, cell signaling and gene expression in cell population averages and in single cells experiments. These results demonstrate how environments that change over time can change key biological processes distinct from traditional approaches. Our results underscore the importance of experimental design to generate precise environmental perturbations that differ from rapidly changing environments. This simple yet general methodology to precisely perturb the cells will enable new understanding and insights into biological processes.

## Methods

### Human Cell Culture and Materials

THP1 (ATCC® TIB-202™) cells were cultured at 0.5-1x 10^6^ cells/ml in RPMI 1640 media (Corning, Catalog#: 15-040-CV) containing 10% Heat inactivated FBS (Gibco, Catalog#: 16140-071), 100 U/ml Penicillin-Streptomycin (Gibco, Catalog#: 15140-122), 2 mM L-alanyl-L-glutamine dipeptide (GlutaMAX™, Gibco, Catalog#: 35050-061) and 0.05 mM 2-Mercaptoethanol (Sigma, Catalog#: M3148) at 37 °C in a 5% CO2 humidity controlled environment.

### Yeast strain and cell culture

*Saccharomyces cerevisiae* BY4741 (MATa; his3Δ1; leu2Δ0; met15Δ0; ura3Δ0) was used for time-lapse microscopy and FISH experiments. To assay the nuclear enrichment of Hog1 in single cells over time in response to osmotic stress, a yellow-fluorescent protein (YFP) was tagged to the C-terminus of endogenous Hog1 in BY4741 cells through homologous DNA recombination. Three days before the experiment, yeast cells from a stock of cells stored at 80 °C were streaked out on a complete synthetic media plate (CSM, Formedia, UK). The day before the experiment, a single colony from the CSM plate was picked and inoculated in 5ml CSM medium (pre-culture). After 6-12 hours, the optical density (OD) of the pre-culture was measured and diluted into new CSM medium to reach an OD of 0.5 the next morning.

### Experimental procedure for human cells upon step and linear gradient stimuli application

A programmable pump (New Era Syringe Pump Systems, NE-1200) was used to apply gradually increasing (linear gradients) profiles. In brief, pumping rate and dispensed volume per interval were calculated as described in the pump profile calculation section and uploaded to the pump via a computer. A syringe pump driving a syringe (BD™, Catalog#: 309628) filled with 5 M NaCl (Corning, Catalog#: 46-032-CV) solution was connected to a needle (Jensen Global, Catalog#: JG21-1.0x) with tubing (Scientific Commodities, Catalog#: BB31695-PE/4). The tubing was inserted into a foam stopper on an autoclaved glass flask (Pyrex, Catalog#: 4980-500) holding the suspension cells. To ensure proper mixing cells were shaken at 100 rpm during the entire experiment using a CO2 resistant shaker (Thermo Fisher Scientific, Catalog#: 88881101). For step stimulation, appropriate amount of 5 M NaCl solution was added by a syringe within 5 seconds to reach the desired final concentration. 5 ml of cells were removed with a syringe (BD™, Catalog#: 309628) through autoclaved silicone tubing (Thermo Scientific, Catalog#: 8600-0020) to collect time point samples. Cell are sampled at 2, 5, 10, 15, 20, 30, 35, 40, 45, 50, 60, 65, 70, 80, 90, 120, and 150 minutes after the start of the experiment.

### Flow Cytometry

Cells were fixed with 1.6% formaldehyde (Fisher, Catalog#: F79-4) in a 15 ml falcon tube. Fixation was quenched by adding 200 mM Glycine after 8 minutes. Cells were washed with PBS (Corning, Catalog#: 46-013-CM) and permeabilized with Methanol (Fisher, Catalog#: A454-4) for 15 minutes on ice. Cells were washed with PBS and blocked with 5% BSA (Rpi, Catalog#: A30075-100.0) in PBS. Cells were then washed and stained with a primary monoclonal antibody conjugated to Alexa Fluor 647 recognizing phosphorylated JNK (Cell Signaling Technologies, Cat.#: 9257) overnight at 4 °C. Flow cytometry was performed on BD LSRII (five lasers).

### Flow Cytometry Analysis

Flow Cytometry data was analyzed with custom R software. The main cell population was gated on FSC-A vs. SSC-A by using the flowcore package^32^. Means and standard deviations of mean fluorescent intensity were calculated for 2-3 replicates and plotted over time.

### Time-lapse microscopy

1.5 ml of yeast cells (Hog1-YFP) in log-phase growth (OD=0.5) were pelleted, re-suspended in 20 µl CSM medium and loaded into a flow chamber. The flow chamber consist of an 1/8” Clear Acrylic Slide with three holes (Grace Bio-labs, 44562), a 0.17µm thick T-shape SecureSeal Adhesive spacer (Grace Bio-labs, 44560, 1L 44040, R&D), a Microscope Cover Glasse (Fisher Sci., 12545E, No 1 22 x 50 mm) and Micro medical tubing (Scientific Commodities Inc., BB31695-PE/4, 100 feet, 0.72mm ID x 1.22mm OD). The cover glass is coated with 0.1mg/ml Concanavalin dissolved in H_2_O. Hyperosmotic perturbations were created using Syringe Pumps as described in the main text (TS experiments) (New Era Pump Systems). On the input of the flow chamber, a three-way valve was connected to switch between a beaker that was used to generate osmolyte concentration profiles and a beaker with media (Beakers 1 and 2, Figure 1c). A syringe pump (Pump 2) at the exit of the flow chamber was used to deliver the specific osmolye profile through the flow chamber with a constant rate. For step-like osmotic perturbation, Beaker 1 had CSM medium with a fixed concentration of 0.4 M NaCl. For the linear gradient perturbation, Beaker 1 only contained media without NaCl at time zero. Using a second syringe pump (Pump 1), profiles of increasing linear osmolyte concentration were generated through pumping 4 M NaCl CSM media into Beaker 1 under constant mixing on a magnetic stir plate (Table S1).

### Image acquisition

The Micro-Manager program (Edelstein et al, 2014) was used to control the microscope (Nikon Eclipse Ti) which is equipped with perfect focus (Nikon), a 100x VC DIC lens (Nikon), a fluorescent filter for YFP (Semrock), an X-cite fluorescent light source (Excelitas) and an Orca Flash 4v2 CMOS camera (Hamamatsu). For step input a pump rate of 0.1 ml/minute was used.

### Single molecule RNA-FISH

Yeast cell culture (BY4741 WT) in log-phase growth (OD = 0.5) was concentrated 10x times (OD = 5) by a glass filter system with a 0.45 μm filter (Millipore). For osmolyte step experiments, cells are exposed to a final osmolyte concentration 0.4 M NaCl, and then fixed in 4 % formaldehyde at specific time points. For linear gradient experiments, concentrated cells are exposed to a linear osmolyte of final concentration of 0.4 M NaCl over 10 minutes, and then fixed in 4 % formaldehyde at specific time points. As described in the main text (TP experiments), a pump was used to inject 4 M NaCl into the beaker with yeast cells (Table S2). Cells are constantly shaken at 250 rpm in a 30 °C incubator to ensure homogenous mixing.

### Fixation, spheroplasting and RNA-FISH probe hybridization

Cells were fixed at 0, 2, 4, 6, 8, 10, 12, 14, 16, 20 minutes after the beginning of applying a linear gradient NaCl of 0.4 M in 10 minutes. Beyond 10 minutes cells were exposed to the fixed 0.4 M NaCl osmolyte. At each time point, 5ml cells are sampled and poured into a 15 ml falcon tube containing formaldehyde resulting in cell fixation at 4% formaldehyde. Cells are fixed at RT for 30 minutes, then transferred to 4 °C and fixed overnight on a shaker. After fixation, cells are centrifuged at 500 × *g* for 5 minutes, and then the liquid phase was discarded. Cell pellets were resuspended in 5 ml ice-cold Buffer B (1.2 M sorbitol, 0.1 M potassium phosphate dibasic, pH 7.5) and centrifuged again. After discarding the liquid phase, yeast cells were resuspended in 1 ml buffer B, and transferred to 1.5 ml centrifugation tubes. Cells were then centrifuged again at 500 × *g* for 5 minutes and the pellet was resuspended in 0.5 ml Spheroplasting Buffer (Buffer B, 0.2% ß-mercaptoethanol, 10 mM Vanadyl-ribonucleoside complex). The OD of each sample was measured and the total cell number for each sample was equalized. 10 μl 2.5 mg/ml Zymolyase (US Biological) was added to each sample on a 4 °C block. Cells were incubated on a rotor for ~20-40 minutes at 30 °C until the cell wall was digested. The cells turn from an opaque to a black color, if they are digested. Cells are monitored under the microscope every 10 minutes after addition of Zymolyase and when ~90 % of the cells turned black, cells are transferred to the 4 °C block to stop Zymolyase activity. Cells are centrifuged for 5 minutes, then the cell pellet was resuspended with 1 ml ice-cold Buffer B and spun down for 5 minutes at 500 × *g*. After discarding liquid phase, the pellet was gently re-suspended with 1 ml of 70 % ethanol and kept at 4 °C for at least 12 hours or stored until needed at 4 °C. Hybridization and microscopy condition were applied as previously described (27).

### Image analysis

Image segmentation was performed in a two-step process. First, the fluorescent tagged Hog1-kinase (Hog1-YFP) image was smoothed, background corrected and automatically thresholded to identify the brightest pixels for each cell. The region of brightest pixels was then used as an intracellular marker. Second, the bright field image was smoothed, background corrected and then overlaid with the previously processed YFP image. On this new image a watershed algorithm was applied to segment and label the objects. After segmentation, objects that are too small, too large or on the border of the image are removed resulting in an image with segmented cells. This process was repeated for each image. After segmentation, the centroid of each cell was computed and stored. Next, the distance between each centroid for each of the two consecutive images was compared. The cells in the two images that have the smallest distance are considered the same cell at two different time points. This whole procedure is repeated for each image resulting in single cell trajectories. For each cell the average per pixel fluorescent intensity of the whole cell (Iw) and of the top 100 brightest fluorescent pixels (It) was recorded as a function of time. In addition, fluorescent signal per pixel of the camera background (Ib) was reported. The Hog1 nuclear enrichment was then calculated as Hog1(t) = [(It(t) − Ib) / (Iw(t) − Ib)]. The single cell traces are smoothed and subtracted by the Hog1(t) signal on the beginning of the experiment. Next, each single cell trace was inspected and cells exhibiting large fluctuations are removed. Bright field images are taken every 10 s with an exposure time of 10 ms and the YFP fluorescent images are taken every 1 minute with an exposure time of 20 ms. During the time points when no fluorescent images are taken, the fluorescent signal from the previous time point was used to segment the cell. Taking images every 10 s with an exposure time of 10 ms using bright field imaging does not result in photo bleaching of the fluorescent signal and ensures better tracking reliability because cells have not moved significantly since the previous image. For cell volume measurements, each time trace was removed of outlier points resulted from segmentation uncertainties. Volume change relative to the volume at the beginning of the experiment was calculated to compare cells of different volumes. For both, the single cell volume and Hog1(t) fluorescent traces, the median and the average median distance were computed to put less weight on sporadic outliers due to the image segmentation process.

## Availability of data and material

The datasets during and/or analyzed during the current study available from the corresponding author on reasonable request.

## Ethics approval and consent to participate

Not applicable

## Consent for publication

Not applicable

## Acknowledgements

AT was funded by an AHA predoctoral fellowship (Award#: 18PRE34050016). HJ, GL and GN were funded by NIH DP2 GM11484901, NIH R01GM115892, and Vanderbilt Startup Funds. The authors thank Benjamin Kesler, Rohit Venkat and Amanda Johnson for comments on the manuscript.

## Authors’ contributions

All authors contributed to the development of the experimental setup. HJ developed the algorithm to compute the pump profiles. GL and AT performed the experiments. AT, HJ and GN wrote the manuscript.

