## Supplementary_Information for "Generating kinetic environments to study dynamic cellular processes in single cells"

*equally contributing

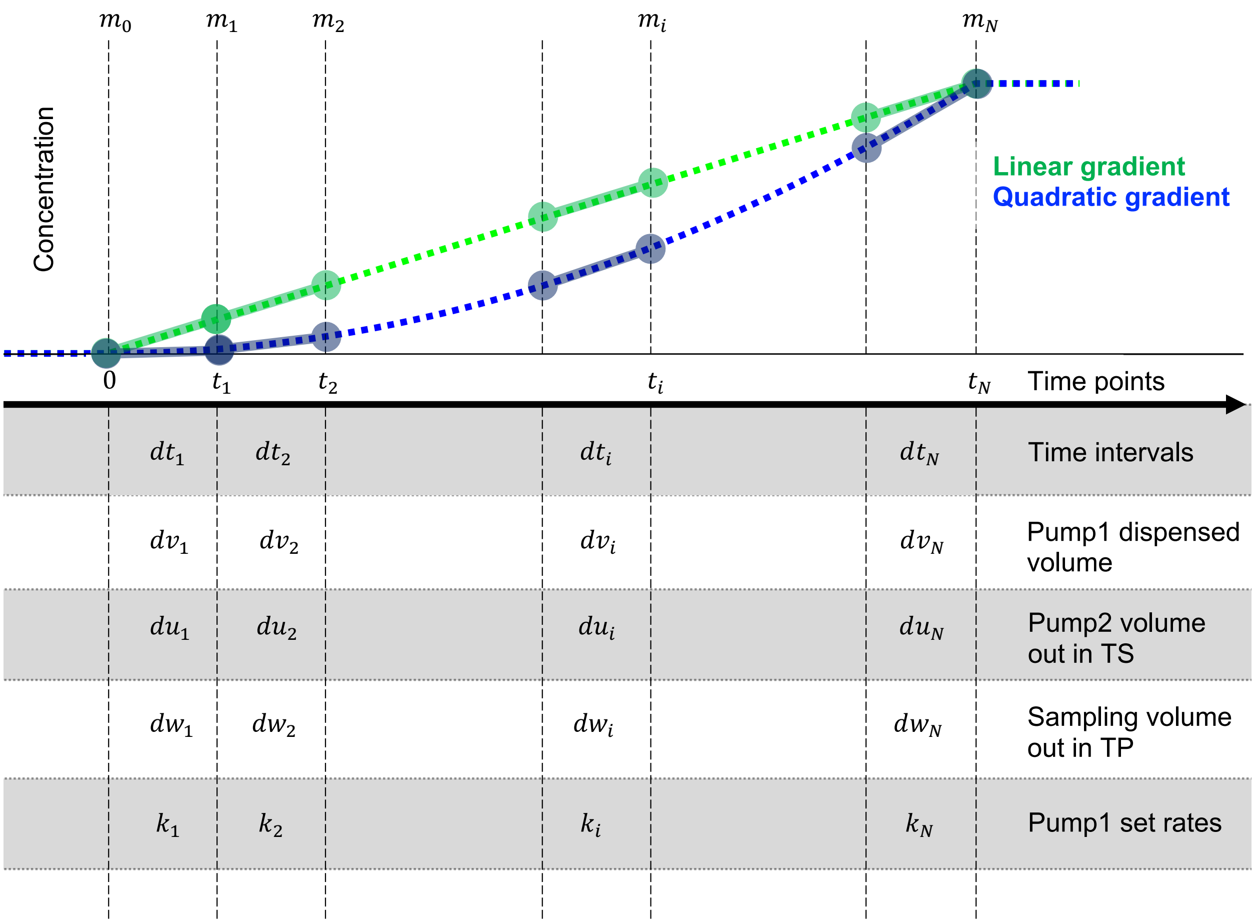


**Figure S1: A diagram illustration of algorithm to compute the pump profiles.** We calculate the stimulus concentration for any profile over discrete time points set by programmable pump by combining several short segments with linear concentration profiles.

**Table S1. Calculation results for TS experiment profile generation.**

| Interval | Time points (min) | Disp. Volume* (mL) | Cumulative Disp. Volume (mL) | Pump rate (µL /min) | Molarity* (M) | Error* % in molarity compared to theory |
|---|---|---|---|---|---|---|
| 1 | 1.25 | 0.125 | 0.125 | 100 | 0.01 | 0 |
| 2 | 2.5 | 0.126 | 0.251 | 100.8 | 0.02 | 0 |
| 3 | 3.75 | 0.126 | 0.377 | 100.8 | 0.03 | 0 |
| 4 | 5 | 0.127 | 0.504 | 101.6 | 0.04 | 0 |
| 5 | 6.25 | 0.127 | 0.631 | 101.6 | 0.05 | 0 |
| 6 | 7.5 | 0.127 | 0.758 | 101.6 | 0.06 | 0 |
| 7 | 8.75 | 0.127 | 0.885 | 101.6 | 0.07 | 0 |
| 8 | 10 | 0.128 | 1.013 | 102.4 | 0.08 | 0 |
| 9 | 11.25 | 0.128 | 1.141 | 102.4 | 0.09 | 0 |
| 10 | 12.5 | 0.128 | 1.269 | 102.4 | 0.1 | 0 |
| 11 | 13.75 | 0.129 | 1.398 | 103.2 | 0.11 | 0 |
| 12 | 15 | 0.129 | 1.527 | 103.2 | 0.12 | 0 |
| 13 | 16.25 | 0.129 | 1.656 | 103.2 | 0.13 | 0 |
| 14 | 17.5 | 0.13 | 1.786 | 104 | 0.14 | 0 |
| 15 | 18.75 | 0.13 | 1.916 | 104 | 0.15 | 0 |
| 16 | 20 | 0.131 | 2.047 | 104.8 | 0.16 | 0 |
| 17 | 21.25 | 0.131 | 2.178 | 104.8 | 0.17 | 0 |
| 18 | 22.5 | 0.131 | 2.309 | 104.8 | 0.18 | 0 |
| 19 | 23.75 | 0.131 | 2.44 | 104.8 | 0.19 | 0 |

| | | | | | | |
|---|---|---|---|---|---|---|
| 20 | 25 | 0.132 | 2.572 | 105.6 | 0.2 | 0 |
| 21 | 26.25 | 0.133 | 2.705 | 106.4 | 0.211 | 0.476 |
| 22 | 27.5 | 0.132 | 2.837 | 105.6 | 0.221 | 0.455 |
| 23 | 28.75 | 0.133 | 2.97 | 106.4 | 0.231 | 0.435 |
| 24 | 30 | 0.134 | 3.104 | 107.2 | 0.241 | 0.417 |
| 25 | 31.25 | 0.134 | 3.238 | 107.2 | 0.251 | 0.4 |
| 26 | 32.5 | 0.134 | 3.372 | 107.2 | 0.261 | 0.385 |
| 27 | 33.75 | 0.134 | 3.506 | 107.2 | 0.271 | 0.37 |
| 28 | 35 | 0.135 | 3.641 | 108 | 0.281 | 0.357 |
| 29 | 36.25 | 0.135 | 3.776 | 108 | 0.291 | 0.345 |
| 30 | 37.5 | 0.136 | 3.912 | 108.8 | 0.301 | 0.333 |
| 31 | 38.75 | 0.136 | 4.048 | 108.8 | 0.311 | 0.323 |
| 32 | 40 | 0.137 | 4.185 | 109.6 | 0.321 | 0.312 |
| 33 | 41.25 | 0.137 | 4.322 | 109.6 | 0.331 | 0.303 |
| 34 | 42.5 | 0.137 | 4.459 | 109.6 | 0.341 | 0.294 |
| 35 | 43.75 | 0.138 | 4.597 | 110.4 | 0.351 | 0.286 |
| 36 | 45 | 0.138 | 4.735 | 110.4 | 0.361 | 0.278 |
| 37 | 46.25 | 0.138 | 4.873 | 110.4 | 0.371 | 0.27 |
| 38 | 47.5 | 0.139 | 5.012 | 111.2 | 0.381 | 0.263 |
| 39 | 48.75 | 0.14 | 5.152 | 112 | 0.391 | 0.256 |
| 40 | 50 | 0.14 | 5.292 | 112 | 0.401 | 0.25 |

**Table S2. Calculation results for TP experiment profile generation.**

| Interval | Time points (min) | Disp. Volume* (mL) | Cumulative Disp. Volume (mL) | Pump rate (µL/min) | Molarity* (M) | Error* % in molarity compared to theory |
|---|---|---|---|---|---|---|
| 1 | 1 | 0.1 | 0.1 | 100 | 0.008 | 0 |
| 2 | 2 | 0.099 | 0.199 | 99 | 0.016 | 0 |
| 3 | 3 | 0.097 | 0.296 | 97 | 0.024 | 0 |
| 4 | 4 | 0.097 | 0.393 | 97 | 0.032 | 0 |
| 5 | 5 | 0.096 | 0.489 | 96 | 0.04 | 0 |
| 6 | 6 | 0.096 | 0.585 | 96 | 0.048 | 0 |
| 7 | 7 | 0.095 | 0.68 | 95 | 0.056 | 0 |
| 8 | 8 | 0.094 | 0.774 | 94 | 0.064 | 0 |
| 9 | 9 | 0.094 | 0.868 | 94 | 0.072 | 0 |
| 10 | 10 | 0.093 | 0.961 | 93 | 0.08 | 0 |
| 11 | 11.667 | 0.154 | 1.115 | 92.4 | 0.093 | 0.357 |
| 12 | 13.333 | 0.154 | 1.269 | 92.4 | 0.107 | 0.312 |
| 13 | 15 | 0.156 | 1.425 | 93.6 | 0.12 | 0 |
| 14 | 16.667 | 0.153 | 1.578 | 91.8 | 0.133 | 0.25 |
| 15 | 18.333 | 0.154 | 1.732 | 92.4 | 0.147 | 0.227 |
| 16 | 20 | 0.156 | 1.888 | 93.6 | 0.16 | 0 |
| 17 | 21.667 | 0.152 | 2.04 | 91.2 | 0.173 | 0.192 |
| 18 | 23.333 | 0.154 | 2.194 | 92.4 | 0.187 | 0.179 |
| 19 | 25 | 0.155 | 2.349 | 93 | 0.2 | 0 |

| | | | | | | |
|---|---|---|---|---|---|---|
| 20 | 26.667 | 0.153 | 2.502 | 91.8 | 0.213 | 0.156 |
| 21 | 28.333 | 0.154 | 2.656 | 92.4 | 0.227 | 0.147 |
| 22 | 30 | 0.155 | 2.811 | 93 | 0.24 | 0 |
| 23 | 31.667 | 0.152 | 2.963 | 91.2 | 0.253 | 0.132 |
| 24 | 33.333 | 0.153 | 3.116 | 91.8 | 0.267 | 0.125 |
| 25 | 35 | 0.155 | 3.271 | 93 | 0.28 | 0 |
| 26 | 36.667 | 0.152 | 3.423 | 91.2 | 0.293 | 0.114 |
| 27 | 38.333 | 0.153 | 3.576 | 91.8 | 0.307 | 0.109 |
| 28 | 40 | 0.154 | 3.73 | 92.4 | 0.32 | 0 |
| 29 | 41.667 | 0.152 | 3.882 | 91.2 | 0.333 | 0.1 |
| 30 | 43.333 | 0.153 | 4.035 | 91.8 | 0.347 | 0.096 |
| 31 | 45 | 0.154 | 4.189 | 92.4 | 0.36 | 0 |
| 32 | 46.667 | 0.151 | 4.34 | 90.6 | 0.373 | 0.089 |
| 33 | 48.333 | 0.153 | 4.493 | 91.8 | 0.387 | 0.086 |
| 34 | 50 | 0.154 | 4.647 | 92.4 | 0.4 | 0 |

*Note on rounding the values: We round the calculated values of $dv_i$ (in mL) to 3 decimal places, which is required by the software of the syringe pump. The resulting calculated pump rates for $k_i$ (in μL /min) are within the range of recommended minimum to maximum pump rate of the syringe pump. The reconstructed molarities and their errors are plotted without rounding in Figure 3, while in Tables 1 and 2 we round them to 3 decimal places for illustration purposes.
